# Sea-ice melt determines seasonal phytoplankton dynamics and delimits the habitat of temperate Atlantic taxa as the Arctic Ocean atlantifies

**DOI:** 10.1101/2023.05.04.539293

**Authors:** Ellen Oldenburg, Ovidiu Popa, Matthias Wietz, Wilken-Jon von Appen, Sinhue Torres-Valdes, Christina Bienhold, Oliver Ebenhöh, Katja Metfies

**Author notes:** **COMPETING INTERESTS** The authors declare no competing interests.

## Abstract

The Arctic Ocean is one of the regions where anthropogenic environmental change is progressing most rapidly and drastically. The impact of rising temperatures and decreasing sea ice on Arctic marine microbial communities is yet not well understood. Microbes form the basis of food webs in the Arctic Ocean, providing energy for larger organisms. Previous studies have shown that Atlantic taxa associated with low light are robust to more polar conditions. In this study, we compared to which extent sea ice melt influences light-associated phytoplankton dynamics and biodiversity over two years at two mooring locations in the Fram Strait. One mooring is deployed in pure Atlantic water, and the second in the intermittently ice-covered Marginal Ice Zone. Time-series analysis of amplicon sequence variants abundance over a two-year period, allowed us to identify communities of co-occurring taxa that exhibit similar patterns throughout the annual cycle. We then examined how alterations in environmental conditions affect the prevalence of species. During high abundance periods of diatoms, polar phytoplankton populations dominated, while temperate taxa were weakly represented. Generally, polar pelagic and ice-associated taxa (such as *Fragilariopsis cylindrus* or *Melosira arctica*) were more prevalent in Atlantic conditions whereas temperate taxa (such as *Odontella aurita* or *Proboscia alata*) have limited potential to persist in colder ice-impacted waters. In contrast to previous assumptions, we think that sea-ice melt acts as a barrier to the horizontal extent of temperate diatoms by preventing their succession at places strongly influenced by polar conditions such as the melting sea ice.

## Introduction

The Arctic is affected by rapid and drastic environmental changes. For instance, air temperatures rise four times (1) as quickly in the region compared to other regions on Earth (2). Arctic sea ice is one of the fastest changing components of the Earth system (3). Over the past decades, the area of Arctic sea ice declined at a rate of about 1 million km^2^ in area extent per decade (3, 4). There are indications for a 40% decline in ice thickness due to thicker and older ice cover (5). The geographical extent of warmer and more saline Atlantic water is expected to expand northwards into the Central Arctic Ocean (CAO), which consequently will become warmer and saltier, further accelerating sea-ice decline (6). This process, called Atlantification of the Arctic Ocean (6), coincides with altered physical conditions. Ecosystems shift towards a more temperate state including the appearance and range expansion of subarctic specie (7, 8, 9, 10, 11, 12, 13). If the temperature increases and the loss of sea-ice continue at their current pace, the Arctic Ocean will likely be seasonally ice-free by 2050 (14). In such a scenario, sea-ice melt-related processes, such as melt-water stratification of the upper layer of the ocean, that is currently observed in the marginal ice zone (MIZ), might become more important over more prolonged periods throughout the seasonal cycle, and a larger geographic area, with ecological consequences for the Arctic Ocean. The MIZ is usually covered with 15- 80% sea ice (15, 16, 17, 18, 19, 20, 21) and its distribution, thickness, and melt dynamics are key drivers of productivity (22), carbon export, biogeochemical cycling, and pelagic-benthic coupling. As a result of decreasing sea ice extent and the expected Atlantification, larger areas of the Arctic Ocean might become favorable for pelagic temperate phytoplankton. As a study site, Fram Strait allows us to investigate the combined effects of Atlantification and seasonal ice cover on Arctic marine ecosystems. Moorings with a suite of physical and biogeochemical sensors, as well as autonomous sampling systems for molecular biodiversity studies (Remote Access Sampler RAS), are positioned at two different locations in Atlantic Waters of Fram Strait at ∼79°N: central Fram Strait (mooring cluster "HG-IV") and in the eastern Fram Strait (mooring cluster "F4") -see Figure 1. F4 is located in the flow path of the West Spitsbergen Current (WSC). HG-IV is located in the vicinity of the interface between the WSC and the East Greenland Current (EGC). The WSC carries relatively warm and salty Atlantic Water via Fram Strait northwards towards the CAO, while the EGC exports cold ice-covered and less saline Polar Water (PW) from the CAO through Fram Strait. In the vicinity of HG-IV, some of the Atlantic Water (AW) is mixed in an eddy-rich area (23) as part of a subduction process (24, 25) with the outflowing colder and fresher water of the EGC. This area is frequently characterized by major sea-ice melt events, as sea-ice coverage regularly extends (26) into the WSC, which carries temperate species towards the CAO. Thus, ecosystem functionality in the vicinity of the MIZ in the WSC might serve as a model for future biodiversity and ecosystem functionality in a seasonally ice-free CAO impacted by Atlantification and thereby inform on the potential of temperate taxa to thrive in a seasonally ice-covered Atlantic-influenced Ocean (27, 28, 29, 30).

**Figure 1:**
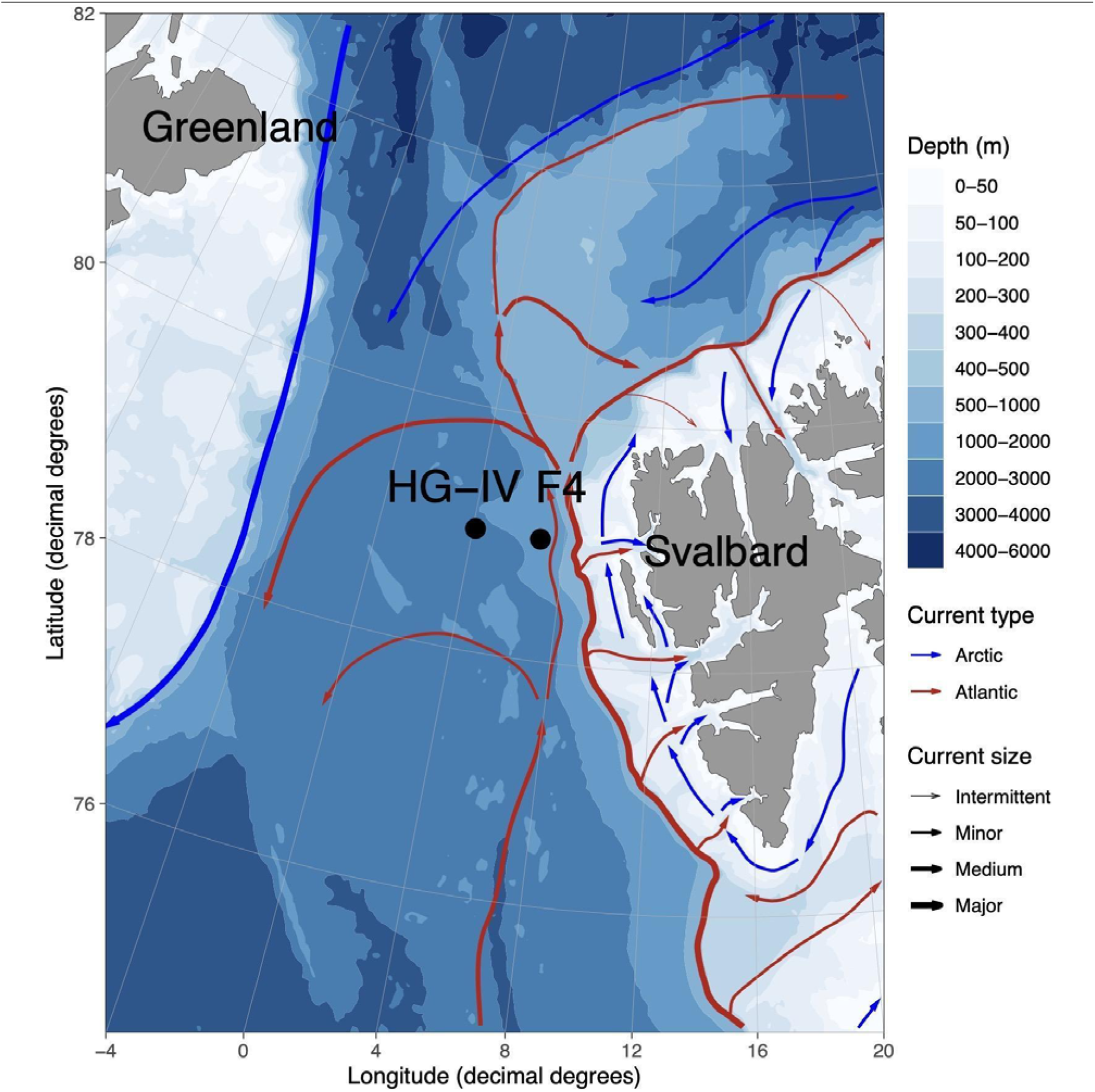
Map of mooring locations, major currents, and water depths in Fram Strait. The main currents in the area are illustrated schematically: West Spitsbergen Current (WSC) in red and East Greenland Current (EGC) in blue. The locations of the moored remote access samplers discussed in this study are marked in black for HG-IV and F4. F4 is located in the WSC and HG- IV west of the WSC. Land is displayed in gray and the different water depths in a white-blue color gradient.

Over the past decades, the transport of sea ice in both volume and velocity towards Fram Strait increased in the area of the Transpolar Drift due to the thinning Arctic pack ice (31, 32, 33). This led to a significant south-eastward extension of the MIZ into Fram Strait during certain years of the past decade. In 2017 the MIZ extended into large parts of the WSC during summer, including the two moorings (33). Conversely, the 2018 ice export was reduced to less than 40% relative to that between 2000 and 2017.

The associated meltwater-induced stratification promoted a longer phytoplankton bloom with a relatively shallow extent and reduced export flux (34). The summer of 2018 had a mixed layer regime and a shorter, more intense bloom compared to other periods. During the spring of that year, there was also an increased carbon export to the deep sea (35). The particularly warm year of 2018 may reflect the conditions of the CAO in the future. The native biodiversity of the communities is a key determinant of whether and how a community or an individual organism can respond to changing abiotic conditions (36). We, therefore, expect that studying the microbial communities and, in particular, comparing the seasonal dynamics between the years 2017 and 2018 can greatly improve our knowledge about the resilience of pelagic and sympagic organisms and how microbial diversity and seasonality scale with the environmental variability. Molecular biodiversity research using ribosomal meta-barcoding has substantially improved our comprehension of marine microbial diversity and distribution patterns during the last 20 years. (37, 38). As part of the FRAM Infrastructure Program (Frontiers in Arctic Marine Monitoring) and the long-term ecological research site LTER HAUSGARTEN, activities in Fram Strait provide information on Arctic marine eukaryotic microbial biodiversity and biogeography based on annually recurring measurements (since 1999) recently expanded by year-round, continuous sampling since 2016. We hypothesize that biodiversity and seasonal succession in the Fram Strait are strongly impacted by sea-ice melt and the extent of stratification (39).

In this study, we exploit this wealth of data through a combination of statistical and bioinformatic approaches. The continuous data collected over two years were decomposed using a Fourier transformation into a series of sinusoidal functions. Each function represents a specific amplicon sequence variant (ASV) dynamic over time. By clustering the ASVs based on their seasonal fluctuation patterns, it became possible to analyze the impact of different water regimes that occurred in 2017 and 2018, as reported in Appen et al. 2021 (34), on both species and community levels. We could elucidate the effects of sea-ice melt on the seasonal dynamics of the associated eukaryotic microbial communities as key drivers of phytoplankton bloom phenology. By assessing the contribution of polar and temperate phytoplankton taxa to eukaryotic microbial communities in the WSC over the annual cycle, we infer the potential of polar taxa to thrive in ice-free Atlantic water and temperate taxa to expand to areas impacted by sea-ice melt.

## MATERIALS AND METHODS

### Sampling

The samples analyzed in this study were collected using McLane Remote Access Samplers (RAS) deployed in conjunction with other oceanographic sensors over three individual annual cycles from June 2016 - August 2019 on long-term moorings at stations HG-IV (79.0118N 4.1666E) and F4 (79.0118N 6.9648E) of the LTER HAUSGARTEN and FRAM in the Fram Strait (40). This study covers the period from January 2017 to December 2018, i.e., two calendar years. One RAS was deployed at a depth between 24-29 m at HG-IV and another at 23-26m - at F4. The RAS samplers contained 48 sterile bags, each collecting water samples of 500 mL at programmed sampling events every two weeks. Samples were preserved by adding 700 µl of half- saturated mercuric chloride (7.5% w/v) to the bags prior to sampling. A sample reflects the pool of up to two samples collected one hour apart in two individual bags. Following the recovery of the RAS devices, water samples were filtered using Sterivex filter cartridges with a pore size of 0.22 µm (Millipore, USA). Filters were then stored at -20°C for later processing.

### Mooring and satellite data

Temperature, salinity, and dissolved oxygen concentration were measured with a CTD-O_2 attached to the RAS frame. Physical oceanography sensors were manufacturer-calibrated and processed as described in (41). Raw and processed mooring data are available at PANGAEA https://doi.org/10.1594/PANGAEA.904565, https://doi.org/10.1594/PANGAEA. 940744, https://doi.pangaea.de/10.1594/PANGAEA.941125. For chemical sensors, raw sensor readouts were used. The fraction of Atlantic and Polar Water were computed for each sampling event following (24) and reported along with distance below the surface (due to mooring blowdown). Sea ice concentration derived from the Advanced Microwave Scanning Radiometer sensor AMSR-2 (42) were downloaded from the Institute of Environmental Physics, University of Bremen (https://seaice.uni-bremen.de/sea-ice-concentration-amsr-eamsr2). Sentinel 3A OLCI chlorophyll surface concentrations were downloaded from https://earth.esa.int/web/sentinel/ sentinel-data-access. For all satellite-derived data, we considered grid points within a radius of 15 km around the moorings. Similar to van Appen et al. 2021 (43), the analyzed datasets consist of ten environmental values for the two locations, F4 and HG-IV, from 01.01.2017 to 31.12.2018. From this dataset, we retrieved the following variables: water temperature (temp °C), fluorescence chlorophyll concentration from in situ sensor (chl_sens ∼*µ*g *l^−^*^1^), daylight (daylight h), water depth (depth m), ice concentration (iceConc %), ice distance (IceDist to 20% ice concentration km), mixed layer depth (MLD m), partial pressure of C*O*_2_ (pC*O*_2__conc *µ*atm), *O*_2_ concentration (O_2__conc *µ*mol *l^−^*^1^), polar-water fraction (PW_frac %).

### DNA-extraction and Illumina amplicon-sequencing of 18S rRNA genes

Isolation of genomic DNA was carried out using the PowerWater kit (Qiagen, Germany) following the manufacturer’s protocol. Obtained DNA was quantified using Quantus (Promega, USA) and stored at -20 °C. 18S rRNA gene fragments from the hypervariable V4 region were amplified by polymerase chain reaction (PCR) with primers 528iF (GCGGTAATTCCAGCTCCAA) and 926iR (ACTTTCGTTCTTGATYRR). illuminaNextV4F (TCGTCGGCA GCGTCAGATGTGTATAAGAGACAGGCGGTAATTCCAGCTCC) and illuminaNextV4R (GTCTCGTGGGCTCG-GAGATGTGTATAAGAGACAGGGCAAATGCTTTCGC) (44). All PCRs had a final volume of 50 µL and contained 0.02 U Phusion Polymerase (Thermo Fisher, Germany), the 10-fold polymerase buffer according to manufacturer’s specification, 0.8 mM each dNTP (Eppendorf, Germany), 0.2 µM *L^−^*^1^ of each primer, and 1 µL of template DNA. PCR amplification was performed in a thermal cycler (Eppendorf, Germany) with an initial denaturation (94 °C, 2min) followed by 35 cycles of denaturation (94 °C, 20 sec), annealing (58 °C, 30 sec), and extension (68 °C, 30 sec) with a single final extension (68 °C, 10 min). The PCR products were purified from an agarose gel 1% [w/v] with the NucleoSpin Gel Kit (Macherey-Nagel, Germany) and Mini Elute PCR Purification kit (Qiagen, Germany). Subsequently, DNA concentrations were determined using a Quantus Fluorometer (Promega, USA). Prior to library preparation, DNA fragments were diluted with TE buffer to a concentration of 0.2 ng µL^-1^. Libraries were prepared according to the 16S Metagenomic Sequencing Library Preparation protocol, and sequenced using MiSeq (Illumina, USA) in 2×300 paired-end runs. Sequence data are available under ENA BioProjects PRJEB43889 and PRJEB43890.

### Sequence analysis

After primer removal using cutadapt (45), reads were processed into amplicon sequence variants (ASVs) using DADA2 v1.14.1 (41), as described in Wietz et al (46). Briefly, reads were trimmed based on quality profiles, with filtering settings truncLen=c(250, 200), maxN=0, minQ=2, maxEE=*c*(3, 3), and trunc*Q* = 0. Followed by merging (minOverlap= 20) and chimera removal, reads were taxonomically classified using PR2 v4.12 (47). The herein reported data has been processed in the scope of autonomous eDNA biodiversity analyses within the FRAM Observatory, as described under https://github.com/matthiaswietz/FRAM-RAS_eDNA.

### Analysis strategy and R packages

All calculations were performed in R version 4.1.3 (2022-03-10). The complete analysis pipeline is available at https://gitlab.com/qtb-hhu/qtb-sda/framstrait_1718. Analysis and plotting tools used for this work are available in a git repository with scripts and an R package. Fourier decomposition was performed with the segmenTier R package (48), available at https://cran.r-project.org/package=segmenTier. The dynamics of eukaryotes were analyzed using the Fourier-transformed time series signals of the relative abundance information. As part of biodiversity, relative species abundance refers to the extent to which a species is common or rare relative to other species in a particular location or community (49). Relative abundance is the percentage composition of an organism of a given species relative to the total number of organisms in that habitat. The data were interpolated on daily bases.

### Time series analysis

Each amplicon sequence variant extracted the time series signal from the relative abundance data using a Fourier approach implemented in the R package segmenTier / segmenTools (50). Fourier transformation is a technique for decomposing functions or signals in the sum of their frequency components, characterized by sine and cosine functions. The Fourier Theorem states that any function can be rewritten as the sum of sinusoidal functions. The approximation becomes more accurate with each additional series element. These elements are called Fourier components.

A measurement for seasonality s for the times series t was calculated by the following formula:

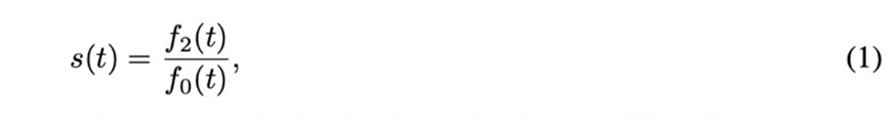

where f_i_ is the i-th fourier component of the times series t and abs is the absolute value function(51, 52).

After the Fourier transformation, the frequency, amplitude, and phase information of each particular ASV time signal was extracted. These values indicate the seasonality, abundance strength, and time of occurrence within the measured period. The choice for the parameter N = 10, the number of clusters for both locations, was chosen to keep the cluster comparable. The metric (Bayesian Information Criterion - BIC) of the applied clustering algorithm proposes a value around 9 and 10 as the optimal cluster number. Groups of species with similar time signals were identified by a clustering approach in the segmenTools R package (50). The significance of overlapping clusters (shared members by two clusters), illustrated as a color gradient, is calculated based on the negative logarithm of the p-value and the number of overlapping features. All identified clusters were classified into low-light, high-light, and mixed-light clusters depending on the light conditions in which their members show the highest abundance. Further, all clusters were named depending on the mooring (H for HG-IV and F for F4) and numbered in ascending order depending on the phase of the sinusoidal function, which was calculated for each cluster from the average of the cluster members. Therefore, the order of the numbers indicates the order of occurrence within the year.

### Conditions preference

To determine population prosperity per year, we calculated the sum of all relative abundances for each ASV within a defined time range. The total abundance values were calculated for F4 and HG-IV locations, respectively. Next, we removed all entries with zero abundance to avoid division by zero and calculated the abundance quotients of 2017 and 2018 and vice versa. Based on the calculated quotients, we defined the preferred condition for each ASV, meltwater regime (MWR), and mixed layer regime (MLR) if the log2(quotient) value is >=1 and accordingly -1. ASV that do not fulfill either condition is assigned to the unspecified group. As a measure of the difference between the locations in a given year for a given group of ASVs, we define the following four quotients:

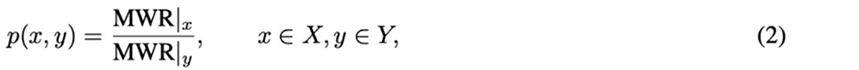

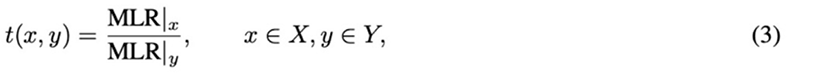

where X = (F417, F418), Y = (HG-IV17, HG-IV18), and MWR (MLR) containing all MWR (MLR) ASVs relative two-year abundances. The restriction is defined by selecting only the AS abundances from the given time and location.

- p(F42017, HG-IV2017) correspond to the ratio of F4 to HG-IV for species preferring the meltwater regime in 2017.
- p(F42018, HG-IV2018) correspond to the ratio of F4 to HG-IV for species preferring the meltwater regime in 2018.
- t(F42017, HG-IV2017) correspond to the ratio of F4 to HG-IV for species preferring the mixed layer regime in 2017.
- t(F42018, HG-IV2018) correspond to the ratio of F4 to HG-IV for species preferring the mixed layer regime in 2018.

To compare how much the meltwater regime is favored on average versus a mixed layer regime within a given site, we define the following equations:

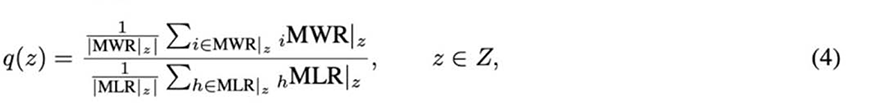

where Z = (F417, F418, HG-IV17, HG-IV18), and MWR (MLR) containing all MWR (MLR) ASVs relative two-year abundances. The restriction is defined by selecting only the AS abundances from the given time and location and _i_MWR (_j_MLR) is the i-th (j-th) relative two-year abundance from the MWR (MLR) ASV.

- q(F42017) corresponds to the ratio for meltwater preference over mixed-layer in 2017 at station F4.
- q(F42018) corresponds to the ratio for meltwater preference over mixed-layer in 2018 at station F4.
- q(HG-IV2017) corresponds to the ratio for meltwater preference over mixed-layer in 2017 at station HG-IV.
- q(HG-IV2018) corresponds to the ratio for meltwater preference over mixed-layer in 2018 at station HG-IV.

### Cross-condition analysis

To investigate how the dynamics of a particular ASV with a preference for a specific water regime change under the conditions of the opposite water regime, we determined and compared the area under the curve (AUC) from the relative abundance within a time range of 365 days. For that, we used on a daily level interpolated abundance data to which we applied a polynomial function and calculated the AUC for each year separately. Afterward, we compared the ratio of the AUC values between the years to illustrate prosperity differences that are related to the environmental conditions of the individual year.

## RESULTS AND DISCUSSION

### Environmental conditions

A pronounced extension of the ice edge/MIZ into the WSC during the first half of 2017, compared to 2018, led to different environmental conditions in this part of the eastern Fram Strait. That mixed layer regime was similar to that expected for a seasonally ice-free Arctic Ocean, impacted by Atlantification. More specifically, eastern Fram Strait experienced extended sea ice melt during spring and early summer 2017. According to van Appen et al. 2021 (34), there were significant differences in environmental conditions between 2017 and 2018, with station HG-IV exhibiting more pronounced differences compared to the pure Atlantic Water station F4. This is best reflected by variability in the fraction of Polar Water, distance to the ice edge, ice concentration, and water column stratification (Figure 2).

**Figure 2:**
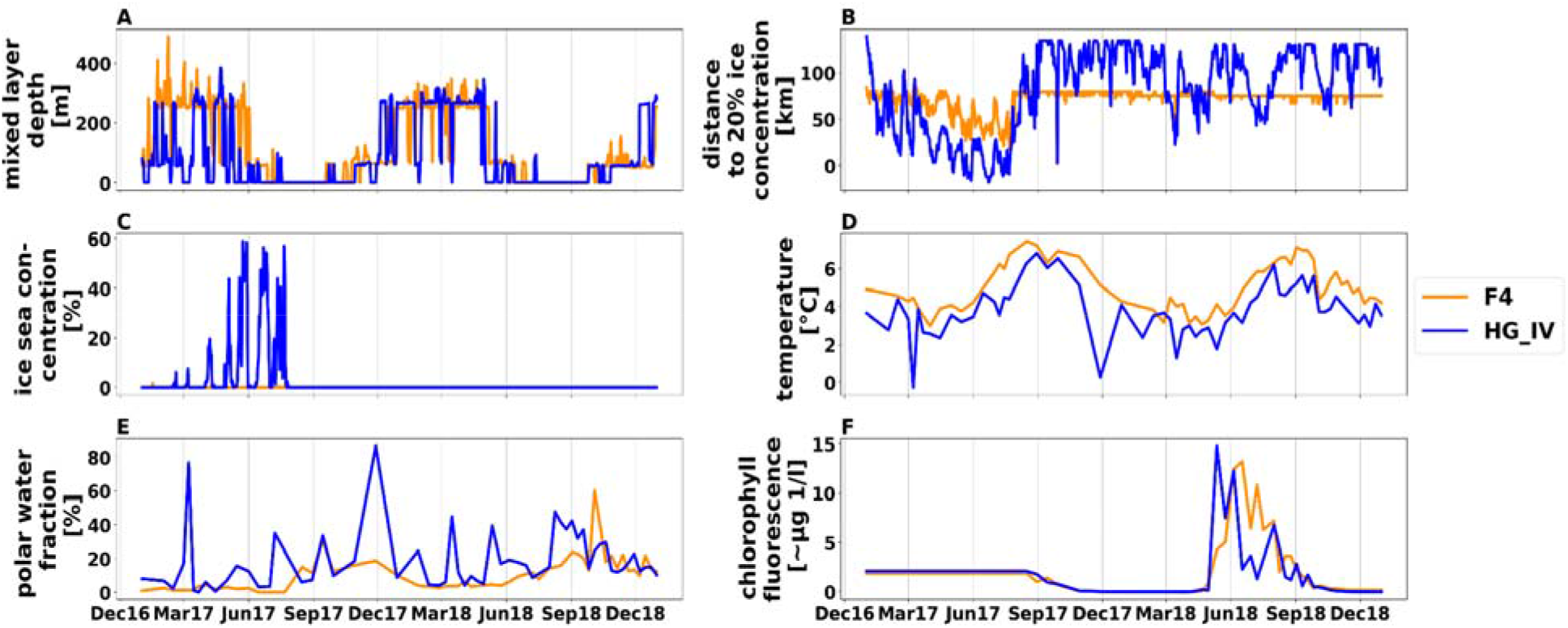
Environmental data for the F4 (dark orange) and HG-IV (blue) location from 2017 to 2018. The x-axis indicates the period from 01.01.2017 to 31.12.2018. The y-axis indicates: **A:** Mixed layer depth (Minimum of the estimated MLD) [m] **B:** Distance to 20% ice concentration (*) [m] **C:** Sea ice concentration [%] **D:** Temperature [◦C] E: Polar water fraction [%] F: Chlorophyll a concentration (**) [µg L−1] *Negative values indicate that the ice edge is south east of the mooring points at the blue curve March 2017 to September 2017) **Sensor did not work before August 2017.

At HG-IV, the mixed layer depth was overall shallower from January to May 2017 compared to 2018 and F4 due to higher ice concentrations. Moreover, HG-IV was frequently impacted by the intrusion of Polar Water (PW) throughout the annual cycle, which is common for this region. Higher fractions of PW were observed for the period’s March, July to August, and November-December of 2017 compared to 2018, according to the RAS data. The intrusion of PW led to lower water temperatures. At HG-IV, temperatures were lower in spring 2017 compared to 2018—ice distances, defined as the distance to 20% ice coverage. At HG-IV, the distance to the ice edge was shorter in 2017 than in 2018 until August but was similar during the remaining months (Figure 2). From mid-August to November; water temperatures were higher in 2017 compared to 2018. In 2017, there was higher ice cover in Fram Strait and subsequent ice melt, resulting in the bloom phenology occurring in a meltwater-stratified water column (MWR). In contrast, in 2018, the bloom phenology occurred in a meltwater-dominated regime (MLR) (34).

At F4, ice distances were not significantly different between the two years. However, water temperatures were higher in 2017 compared to 2018 from mid-August to November.

In the following, we investigated the behavior of eukaryotic microbes under different water regimes (melt water and mixed layer). For that, we used a top-down structure to describe the abundance changes over time for i) all ASVs, ii) specific ASV clusters, and iii) single key species.

### Preference of eukaryotic microbes for the different water regimes

There is a remarkable similarity in ASV composition at both stations. 50% (583) of the ASVs were detected at these two stations, while 22% were unique to F4 (254 ASVs) and 28% to HG-IV (320 ASVs) (Figure S6, Figure S2). Considering the number of ASVs at each station as a baseline, 583 (64.56%) of the 903 ASVs found at HG-IV were also detectable at F4. Conversely, 583 (69.65%) of the 837 ASVs at F4 were present at HG-IV. To analyze the taxa peak abundance of the microbial eukaryotes under different regimes at HG-IV, the total relative abundance for each ASV per year was calculated and compared between the years. Based on that rate, the ASVs were sorted into three groups: the unstratified mixed layer regime (MLR), the highly stratified meltwater regime (MRW), and an unspecified group. The MLR group includes all ASVs (called temperate taxa), which were two times more abundant in HG-IV-2018 compared to HG-IV-2017 (n=67 [11.49% of the shared ASVs]). In contrast, ASVs that were two times more abundant in HG-IV-2017 compared to HG-IV-2018 are members of the MWR group (n=94 [16.12% of the regime shared ASVs]), which were named polar taxa. The remaining ASVs were sorted into an unspecific group (n=422 [72.38% of the regime-shared ASVs]). The last group was excluded from the following analysis. Notably, out of the 583 shared ASVs, only 161 were regime specific in this study, which are distributed between the groups MLR (41.61 %) and MWR (58.38 %) [Table S5].

We compared both groups to identify differences attributed to either location (as shown in Figure 1) or the varying conditions between 2017 and 2018. To do so, we conducted two types of comparisons: i) within each year, we compared the stations to each other, and ii) within each station, we compared the data from 2017 and 2018. First, we compared the relative abundance differences in 2017 between stations. Therefore we calculated the median of the MLR group and MWR group, respectively, for F4 and HG-IV and compared them with each other. Our results showed that the median differences of species favoring mixed-layer were 1.54 times larger than the median differences of the species favoring meltwater in 2017 (Table S5; see methods formula (2,3)). In addition, we could confirm the same observation when comparing the relative abundances of each ASV member of the above groups (one-sided Kolmogorov-Smirnov test p-value: 3.13E-05). In the next step, we repeated the same analysis for the year 2018.

In contrast to 2017, the median differences in 2018 of the meltwater-favoring species were 2.78 times greater than the median differences of the mixed-layer favoring species (Table S5; see methods formula (2,3)). Also, in this case, comparing the relative abundance of the corresponding ASVs could support this observation (one-sided Kolmogorov-Smirnov test p-value: 1.376E-14). Once we had distinguished dissimilarities among the stations, our attention turned to describing dissimilarities over the years. This was motivated by the different water regimes observed in 2017 and 2018 (34). Consequently, this examination enabled us to demonstrate how species abundance is influenced by varying environmental circumstances. Therefore we compared the relative abundance ratio within the same group (MLR, MWR) but between years (2017 vs. 2018). The difference between the two years (2017 and 2018) for each group was less significant at station F4 (MWR=1.23 and MLR=0.60), whereas at HG-IV, the discrepancy was approximately four times higher than that observed at F4 for the same years (MWR=2.13 and MLR=0.27), see Table S5. As a result, we used station F4 as a reference for the constant environment because it is less influenced by meltwater conditions. In contrast, the HG-IV location offers the opportunity to study the effects of Atlantification in a seasonally ice-covered Arctic Ocean, conditions that are expected for the CAO in the near future (53). For that, we examined how each other’s water regimes affected the relative abundance of the respective ASV. We aimed to determine whether polar or temperate ASVs were more resilient to the opposing condition. For the analysis, we specifically selected ASVs that are known to grow in polar or temperate conditions (54, 55, 56, 57, 58, 59, 60, 61).

**Figure 3:**
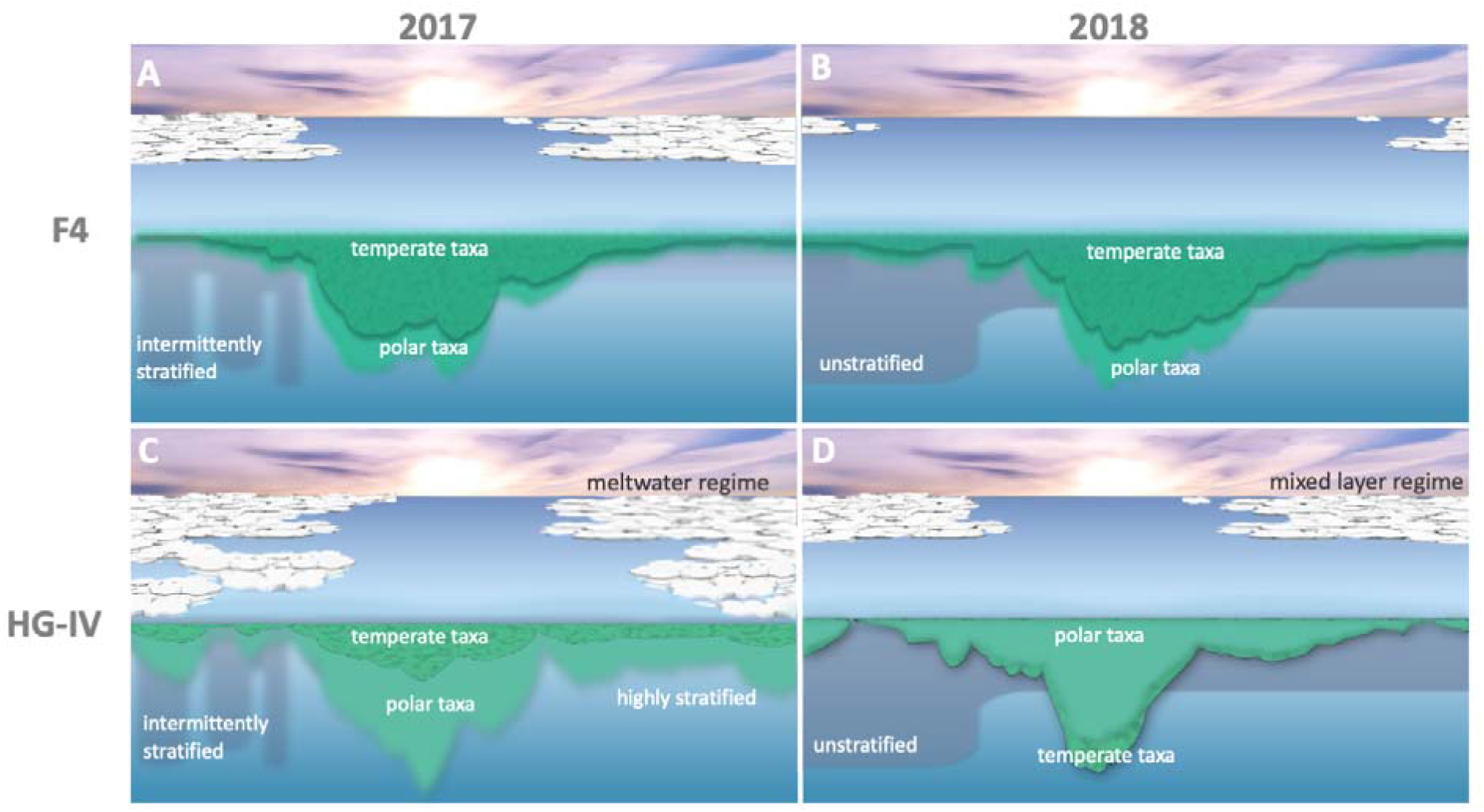
Effects of meltwater and mixed layer conditions on temperate (dark green) and polar (light green) taxa. The x-axis shows the months January through December from 2017 through 2018. The green areas reflect the relative abundances of temperate (dark green) and polar (light green) taxa as extracted from the data. Since the data is relative, no quantification is given on the y-axis. The relative abundance curves of A and B were derived from water column samples from cluster F-06, and C and D from cluster H-06. **A:** Polar and temperate taxa are observed in similar abundances in the highly stratified meltwater regime at F4 in 2017. **B:** Similar abundances for polar and temperate taxa in the mixed layer regime at F4 in 2018. **C:** Reduced abundance of temperate taxa in the meltwater regime with high stratification at HG-IV in 2017. **D:** Reduced abundance of polar taxa in the mixed layer regime at HG-IV in 2018.

### Seasonal succession of eukaryotic microbes

To understand the seasonal succession of eukaryotic microbes, we analyzed the phases obtained from the sinusoidal function after Fourier transformation. This allows us to determine the chronological timeline of the species in this region. Community detection from the time series analysis of 837 and 903 ASVs from the F4 and HG-IV moorings revealed ten clusters of seasonally concerted and ordered occurrences of eukaryotic microbial species (Figure 4, Table 1) throughout the observation period. The frequency obtained from the sinusoidal function (light grey) shows the number of high abundance periods of each community per year. Most clusters (85%) had two maxima, indicating that most organisms exhibit a seasonal occurrence with the highest abundance once a year (Figure 4, Table 1). We divided the clusters based on their high abundance period into two classes of light conditions. The low-light (LL 0-2 hours sunlight per day) clusters include species with a high abundance phase in the low-light period from October to March when water temperature and distance to the ice edge are low. The high-light class (HL 2-24 hours sunlight per day) includes clusters, in which the high abundance phases coincide with the high-light period from March to October. All other clusters are collected in the mixed light (NA) class. This distinction allowed us to test the succession of the organisms regarding environmental factors per light condition separately. Next, we compared the species distribution between the two moorings in terms of abundance and seasonality to first test for commonalities and differences between both sampling sites and second, to measure the succession and prosperity of common species regarding the different water regimes. In addition, we compared the time series cluster composition from HG-IV and F4 with each other to identify overlapping communities between both locations. For example, the similarity in cluster composition between the two moorings was highest during the high-light period, particularly between clusters H-06 and F-06 and clusters H-08 and F-08 (Figure 5). The presence of these common ASVs at both mooring sites can be explained by a similar trend in the transportation of temperate organisms through the northward-flowing warmer Atlantic and the transportation of polar organisms through the intrusion of polar water from EGC. This pattern was also observed for zooplankton (62, 63). On the other hand, the varying quantities of ASVs reaching each station because of variations in the influence of the two currents may also explain the biodiversity observed at these two locations (Figure S1).

**Figure 4:**
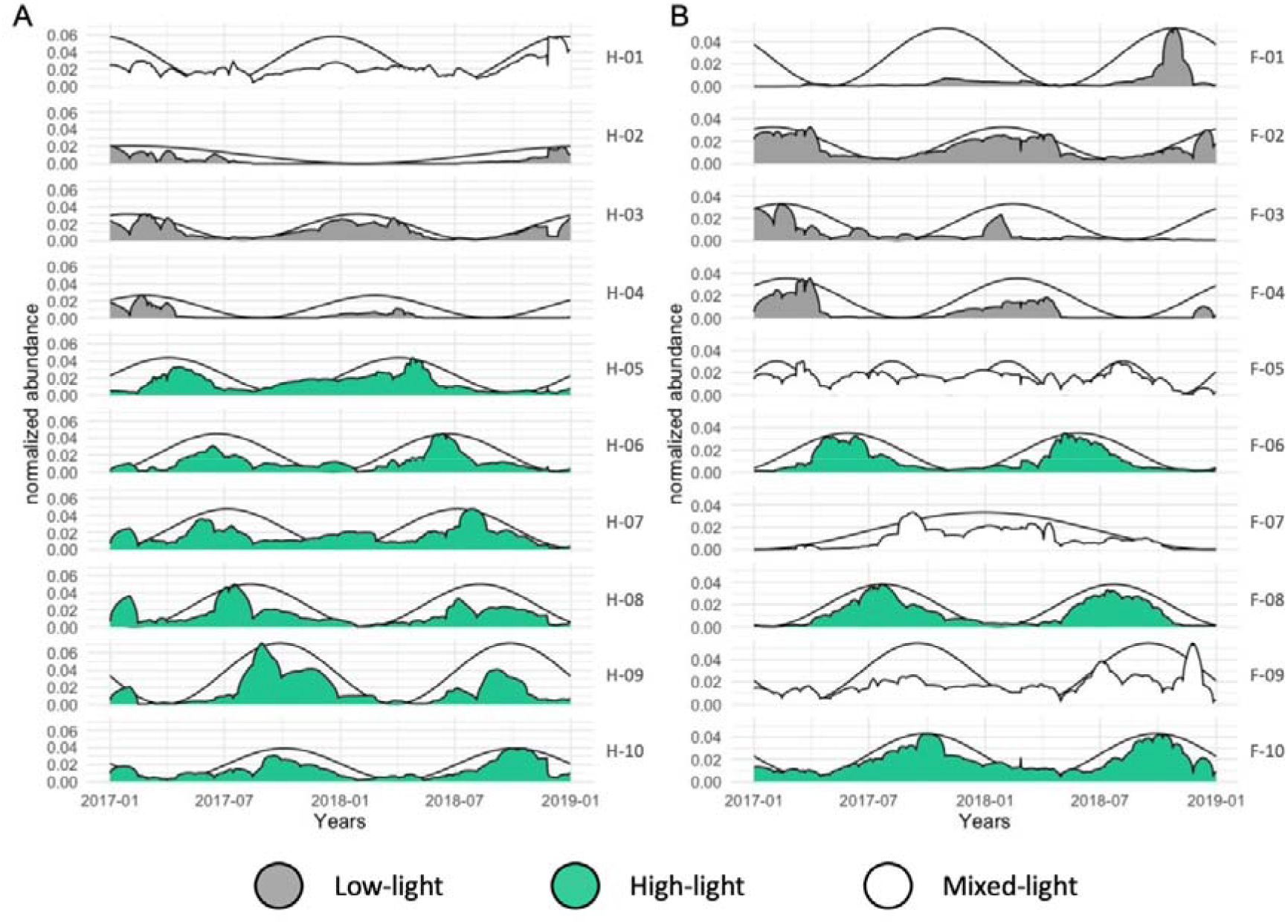
Time-Series Clustering for both moorings spanning the years 2017-2018. The x-axis indicates the period from 01.01.2017 to 31.12.2018. Black sinusoidal curves show the predicted seasonality of the entire cluster based on the dominant Fourier component. The respective relative abundance is shown for each cluster on the left y-axis. Cluster names are shown on the right. The clusters are sorted by phase which illustrates the time of maximal abundance of each community. Clusters are colored according to the three classes HL (green), LL (grey), and NA (white) introduced in the text. **A:** HG-IV, **B:** F4

**Figure 5:**
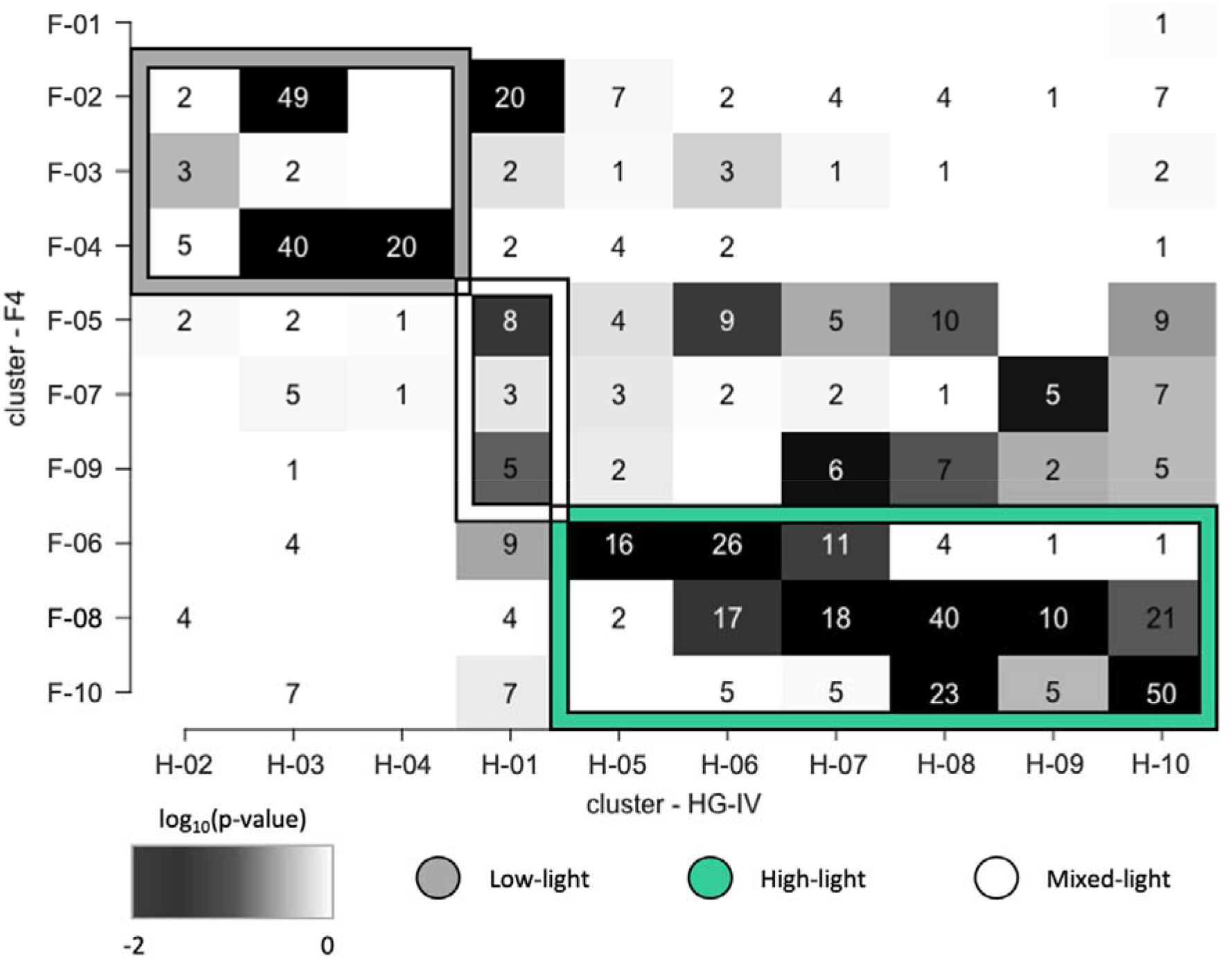
Cluster overlap between F4 and HG-IV locations. The clusters of F4 are plotted on the y-axis against the clusters of HG-IV. The numbers inside the boxes indicate how many ASVs are shared between two clusters The clusters of each location are sorted according to their classes: low-light (grey box frame), mix-light (white box frame) and high-light (green box frame) from top to bottom (F4) and from left to right (HG-IV). The background color of the boxes shows the significance of the overlap from dark (highly significant) to white (non significant).

**Table 1.** Cluster Overview with the ten clusters for the moorings F4 and HG-IV. The cluster names, light types (high-light (HL), low-light (LL), mixed-light (NA)), the number of peaks and the total cluster size of ASV and the percent size, the s-score that measures the seasonality, the area under the curve (AUC) for both years (see methods), the. quotients of those years and the number of ASV that only occur on this mooring: absolute (MS(ab.s)) and relative values (in %) (MS(rel)) (MS: mooring specific).

| name | type | #peaks | cl_size | cl_size % | s-score | AUC17 | AUC18 | AUC17/18 | AUC18/17 | MS(abs) | MS(rel) |
| --- | --- | --- | --- | --- | --- | --- | --- | --- | --- | --- | --- |
| H-01 | NA | 2 | 72 | 9 | 0.18 | 6.9837 | 8.8812 | 0.7863 | 1.2717 | 12 | 16.67 |
| H-02 | LL | 1 | 68 | 8 | 0.39 | 1.81 | 1.0084 | 1.7949 | 0.5571 | 52 | 76.47 |
| H-03 | LL | 2 | 151 | 18 | 0.44 | 4.4387 | 4.1164 | 1.0783 | 0.9274 | 41 | 27.15 |
| H-04 | LL | 2 | 50 | 6 | 0.83 | 1.7481 | 0.7188 | 2.432 | 0.4112 | 28 | 56 |
| H-05 | HL | 2 | 76 | 9 | 0.33 | 5.2585 | 5.1262 | 1.0258 | 0.9748 | 37 | 48.68 |
| H-06 | HL | 2 | 87 | 10 | 0.41 | 4.1827 | 4.7476 | 0.881 | 1.1351 | 21 | 24.14 |
| H-07 | HL | 2 | 65 | 8 | 0.2 | 6.0155 | 6.5539 | 0.9179 | 1.0895 | 13 | 20 |
| H-08 | HL | 2 | 113 | 14 | 0.32 | 6.6398 | 4.405 | 1.5073 | 0.6634 | 23 | 20.35 |
| H-09 | HL | 2 | 33 | 4 | 0.53 | 7.9501 | 4.5641 | 1.7419 | 0.5741 | 9 | 27.27 |
| H-10 | HL | 2 | 122 | 15 | 0.41 | 5.0174 | 5.0116 | 1.0012 | 0.9988 | 18 | 14.75 |
| F-01 | LL | 2 | 27 | 3 | 0.66 | 0.6775 | 2.8243 | 0.2399 | 4.1687 | 26 | 96.3 |
| F-02 | LL | 2 | 144 | 16 | 0.39 | 5.4081 | 5.1656 | 1.0469 | 0.9552 | 48 | 33.33 |
| F-03 | LL | 2 | 33 | 4 | 0.46 | 3.289 | 1.228 | 2.6783 | 0.3734 | 18 | 54.55 |
| F-04 | LL | 2 | 168 | 19 | 0.75 | 2.9621 | 1.8344 | 1.6148 | 0.6193 | 94 | 55.95 |
| F-05 | NA | 1 | 61 | 7 | 0.06 | 6.194 | 5.2571 | 1.1782 | 0.8487 | 11 | 18.03 |
| F-06 | HL | 2 | 109 | 12 | 0.58 | 3.9653 | 4.1903 | 0.9463 | 1.0567 | 37 | 33.94 |
| F-07 | NA | 4 | 53 | 6 | 0.13 | 3.4143 | 3.529 | 0.9675 | 1.0336 | 24 | 45.28 |
| F-08 | HL | 2 | 142 | 16 | 0.58 | 4.8684 | 4.4017 | 1.106 | 0.9041 | 26 | 18.31 |
| F-09 | NA | 2 | 36 | 4 | 0.17 | 5.6129 | 7.6206 | 0.7365 | 1.3577 | 8 | 22.22 |
| F-10 | HL | 2 | 130 | 12 | 0.34 | 7.0415 | 7.0604 | 0.9973 | 1.0027 | 28 | 21.54 |

### Low-light period

From October to March, the low-light period was characterized by the ordered appearance of four clusters at each mooring (F4: F-01, F-02, F-03, F-04; HG-IV: H-02, H-03, H-04). At both stations, these clusters contained *∼*40% of the total ASVs, and *∼*50 % of the total reads. The clusters were dominated by heterotrophic dinoflagellates, parasitic Syndiniales, and other small heterotrophic flagellates like MAST and Picozoa (Figure S4) This mainly heterotrophic community composition resembles prior reports of microbial diversity for the low light period in the Arctic Ocean (63, 64, 65) possibly linked to feeding on bacteria (64). However, in all low-light clusters a set of diatom ASVs are present, which possess considerable relative abundances at both stations (Figure S4). The relative sequence abundances of these ASVs were higher at HG-IV than at F4. These ASVs are ice-associated genera or species, such as *Melosira arctica*, *Navicuales sp.,* or *Attheya sepentrionalis* (Figure S4 Table S4). Members of these taxa are adapted to low light conditions and colder temperatures (66). They usually live in or under the ice (67). Thus physical exchange processes at the interface between water and sea ice and advection might have been the sources of these diatoms in the water column during winter at HG-IV. The persistence of diatoms (Bacillariophyceae) during the polar night in ice-covered waters was previously observed in the CAO (64) and in a year-round molecular study of eukaryotic microbes in Isfjorden (West Spitsbergen) (65). However, their survival strategies and ecological roles during winter remain primarily unresolved (65). Resting stages such as spores or cysts are a potential strategy of Bacillariophyceae and Dinophyceae to persist in unfavorable conditions like the Arctic winter (68). The taxon-specific survival of diatoms during winter in and under the ice is thought to drive the composition of Arctic phytoplankton during early spring. Diatoms maintain chlorophyll in their cells during the polar night, which gives them a growth advantage at the time of light return (69), when diatoms are the major primary producers in Arctic marine ecosystems.

### High-light period

The high-light period (March to October) was characterized by a consecutive appearance of three clusters at F4 (F-06, F-08, F-10) and six clusters at HG-IV (H-05, H-06, H-07, H-08, H-09, H-10), respectively (Table 1). The high-light clusters contained ∼50 % of all mooring specific ASVs (Table 1). The member composition of the earlier high-light clusters H-05, H-06, and H-07 in 2017 was similar to the composition of the earlier high-light clusters F-06, F-08, and F-10 in 2018 (Figure 5). This suggests that both stations shared a similar community at the beginning of the high-light period (Table S3, Table S4). During this period, sequences of diatoms and other autotrophic taxa, either dominated or were highly abundant besides dinoflagellates (Figure S4, Table S3). Regarding the order of the sequential appearance of the diatoms over the year, we first compared the clusters with ASVs that showed an increased abundance during spring (H-05 and F-06). At HG-IV, diatom ASVs were largely affiliated with the Arctic diatoms *Fragilariopsis cylindrus, Bacillaria paxilifer, Chaetoceros neogracilis* and *Grammonema striatula* (70, 71, 72) (Table S3) Their major contribution to the pelagic spring bloom was in line with previous observations (28, 73, 74) emphasizing the polar character of the spring bloom community at HG-IV. The polar taxa *Grammononema striatula* and *Chaetoceros neogracilis* were also highly abundant in cluster F-06 (constituting 22% of the total cluster abundance), the first high-light cluster of station F4. In contrast, the temperate taxon *Odontella aurita (75)* was among the five most abundant diatoms (constituting 6% Table S4 F-06) observed in this cluster. The presence of *Odontella aurita* illustrates the influence of Atlantic Water and the concurrent advection of organisms from more temperate waters at this station (Figure 6). *Odontella aurita* is also known to be a major contributor to spring blooms in the German Bight (76).

**Figure 6:**
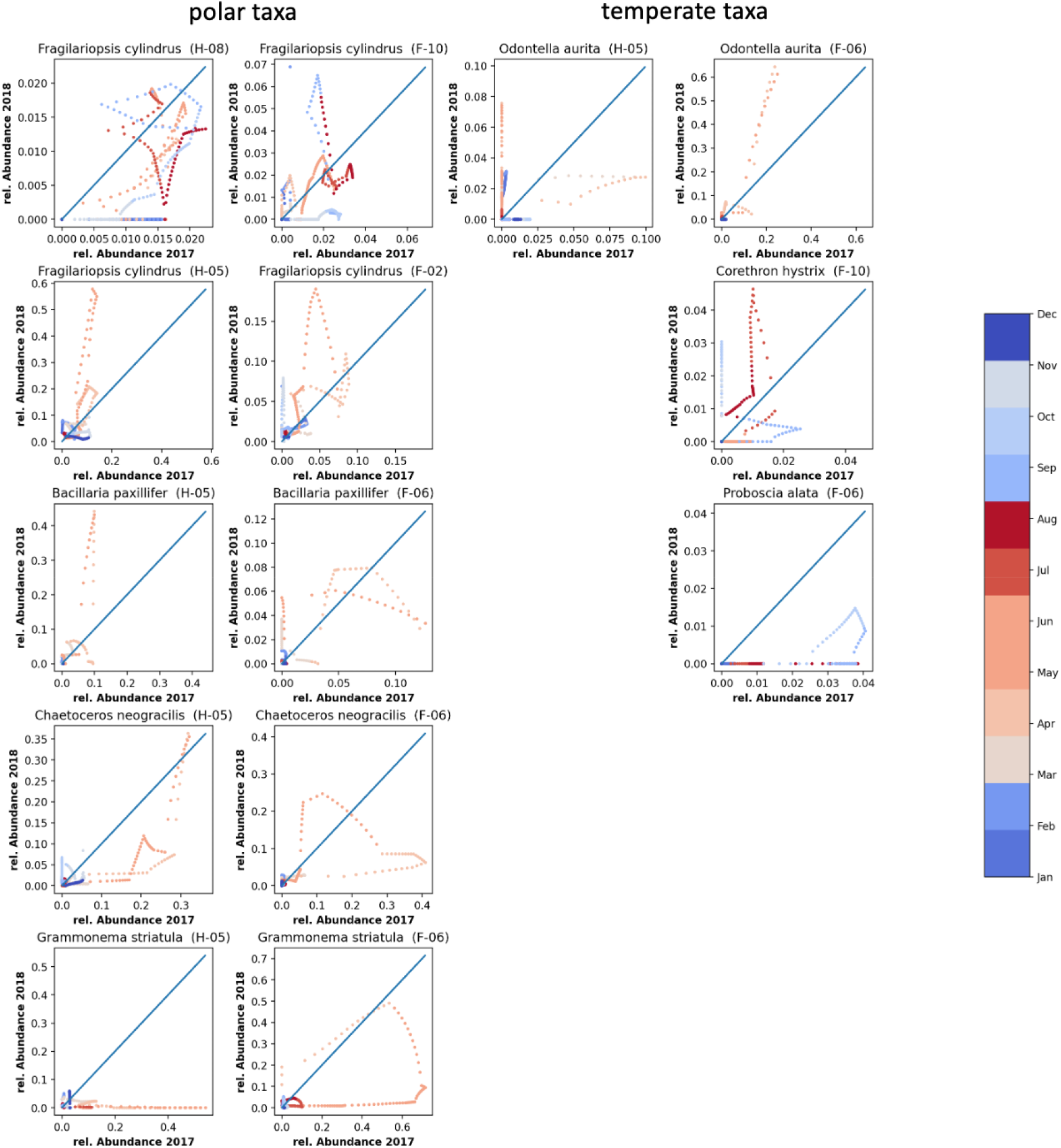
Correlation between the relative abundances of selected ASVs in 2017 vs 2018. The diagonal (blue line) indicates the line on which abundances in 2017 and 2018 would be identical. On the left side (first and second columns) selected polar taxa are displayed, where the first column shows the species at HG-IV and the second column the same ASV at F4. The right side shows selected temperate taxa, where the third column displays species at HG-IV and the fourth column the same ASV at F4. The dots indicate: *Fragilariopsis cylindrus* (ASV207: H-08, F-10), *Fragilariopsis cylindrus* (ASV16: H-05, F-02), *Bacillaria paxillifer* (ASV98: H-03, F-06), *Chaetoceros neograciis* (ASV17: H-05, F-06), *Grammonema stratula* (ASV33: H-05, F-06), *Odontella aurita* (ASV96:H-05, F- 06), *Corethron hystrix* (ASV172: F-10), *Proboscia alata* (ASV947: F-06), Color bar and colored dots indicate month of the year from blue (winter) to red (summer).

Differences in diatoms composition between F4 and HG-IV were even more pronounced in the late summer clusters (H-08 and F-10), having their highest abundance after July. At HG-IV, cluster H-08 was dominated by sea-ice-associated diatoms such as *Melosira arctica* and other ice-related taxa (67) such as *Fragillariopsis sublineata* and *Fragillariopsis cylindrus*, *Chaetoceros rostratus*, or *Thalassiosira sp.* which contribute with 57% (Table S4 H-08) of the total diatom abundance. In contrast, the diatom community in the late cluster F-10 was dominated by *Pseudonitzschia sp* (contributing 38% of the total diatom abundance Table S4 F-10), while the polar taxa that dominated cluster H-08 were only present with smaller contributions. Moreover, this cluster contained significant amounts of *Corethron hystrix* and *Proboscia alata* (2% of the total abundance of the diatoms Table S4 F-10, Top11). Those two species thrive in temperate waters (77, 78) illustrating the impact of Atlantic advection at station F4. In cluster H-09, which accounted for 28% of the total abundance in Table S3, the genus Pseudonitzschia was the dominant species. Additionally, in the second cluster at HG-IV (H-02), Pseudonitzschia contributed to a bloom occurring later in the year, specifically in the autumn season (79). Studies have shown that this diatom undergoes blooming throughout the year, typically exhibiting a minor bloom in June, followed by a more substantial bloom in late August or early September (79). Peak sequence abundances of other major Arctic pelagic autotrophs such as *Phaeocystis sp.*, *Chaetoceros socialis* or *Micromonas sp.* were mainly restricted to high-light clusters (Table S3). Notably, sequence contributions represented by ASVs of *Phaeocystis pouchetii* were highest in the early spring clusters (H-05 & F-06) accounting for 16 % and 11% of the total abundance respectively (Table S3 H-05, F-06). This result agrees with previous observations in the WSC and under the ice north of Svalbard (80, 81). Nonetheless, *Phaeocystis sp.* and *Micromonas sp.* were found even during winter. Their relative contributions to the eukaryotic microbial community remained below levels observed in other molecular genetic studies of the deep chlorophyll maximum (DCM) in the Fram Strait during summer (82). Although there may be ecological reasons for the under-representation of small taxa in this study, the possibility that RAS was biased towards larger eukaryotic microbial cells can not be ruled out.

### Impact of sea-ice melt on seasonal phytoplankton dynamics and consequences for bloom phenology in Atlantic waters

The different environmental conditions observed in 2017 and 2018 did not seem to affect the order of the annually recurring community clusters at F4 and HG-IV. Instead, changes in environmental conditions resulted in differences in their persistence, abundance amplitude, and integrated abundances (Figure 4A, Figure 4B). At F4, environmental conditions during the high light periods of 2017 and 2018 were similar. In consequence, the integrated seasonal cluster abundance, reflected by the area under the curve, did not significantly change from one year to the other (Table 1). In contrast, we observed differences between both years for HL and LL periods at HG-IV (Figure S5). According to our data, the changes in environmental conditions, associated with sea-ice melt in spring and summer 2017 at HG-IV, might have significantly affected the communities during high-light periods. For example, these changes can be observed in the high-light cluster H-09 (Table 1). The last period of the cluster (2018) shows a 1.7-fold decrease in abundance compared to the first period (2017). Despite the area under the curve of the early high-light clusters (H-05, H-06, H-07) showing almost no difference between the two years at HG-IV, the amplitude was much lower in 2017 compared to 2018 (Table S2). This observation suggests that the growth rates in 2017 were lower. It is important to note that the organism abundances only reflect relative proportions of the filtered samples. However, in 2017 the RAS was below the productive layer for at least the first half of the high-light period (37), which may explain the lower relative abundances.

Polar pelagic taxa, such as *Chaetoceros neogracilis* and *Grammonema striatula*, were dominant (compared to other Bacillariophyta) in the first clusters of the high-light period at both stations (H-05, F-06, Table S3). These species are more robust to variation in ice coverage. In contrast, the contribution of *Fragillariopsis cylindrus* to the spring cluster H-05 was greater at HG-IV than at F-06, as indicated in Table S3. During the spring of 2017 at HG-IV, lower relative abundances of *Fragilariopsis cylindrus* may suggest lower growth rates, which could be attributed to higher ice coverage at this station. *Fragilariopsis cylindrus* and *Bacillaria pacillifer* were among the ASVs with the ten highest relative abundances at both stations. They had higher relative abundances during the spring at HG-IV compared to F4 in the observation period, as shown in Table S4 and Figure 6. This was likely because they benefited from lower ice concentrations and comparatively higher water temperatures at HG-IV during the spring of 2018 compared to 2017 (Figure 6). This observation suggests that these polar taxa are not strictly dependent on polar conditions and can tolerate or benefit from Atlantic influence.

*Odontella aurita*, a temperate taxon occurring at both stations, benefits at both stations from warmer temperatures. The contribution of this temperate species in cluster H-05 was negligible, accounting for only 1% of total abundance, with a further decrease in 2017 to 0.81%, indicating that it struggles to thrive under the ice. In contrast, at mooring F4, its contribution to the spring cluster F-06 was high in both years (Table S3) as temperatures were in a similar range. During the later part of the season in HG-IV, the area under the curve of cluster H-08 showed a 1.5-fold increase in 2017 compared to 2018, as indicated in Table 1. This cluster mainly comprised typical sea-ice-associated diatoms like *Melosira arctica*, *Fragillariopsis sublineata* and -*cylindrus*, and *Chaetoceros rostratus*. Interestingly, these diatoms did not contribute significantly to the phytoplankton community at F4 during the same time of the year. This indicates a sea-ice melt-related release of sea-ice-associated taxa. The environmental conditions existing at this time, especially meltwater stratification, promoted their bloom in the Atlantic Water of Fram Strait (Table S3 Table S4).

During the specified time frame, there was a notable decrease in the prevalence of polar spring phytoplankton species at the start of the season, accompanied by a corresponding increase in the abundance of ice-associated phytoplankton species during the autumn of 2017. It is worth noting that the peak abundance of ice-associated phytoplankton species usually occurs later in the season in the CAO (83, 84, 85). Ice-associated phytoplankton is less present at HG-IV in 2018 (ice-free year) and does not significantly contribute to the autumn community at ice-free station F4 in either year.

## CONCLUSION

In this study, we compared the dynamics of phytoplankton ASVs from two locations in the Fram Strait (moorings HG-IV and F4) as recorded in 2017 and 2018. Although data from only two years are not necessarily representative of the long-term development of environmental parameters, these particular years exhibit conditions that make them appear ideal for comparing current conditions with those expected in the future in an Atlantified CAO. This comparison supports a new perspective on how the eukaryotic microbial community in the Central Arctic Ocean might change in the near future. Climate change will likely lead to an ice-free Central Arctic Ocean in summer but ice-covered in winter, as suggested by some climate model scenarios (14).

In our analysis, we could show that a meltwater regime can strongly influence arctic micro-eukaryotes on several levels and that phytoplankton bloom phenology in 2017 is a result of increased sea ice melt (34). We could extend previous observations about the influence of sea-ice melt on community dynamics and carbon export. We propose that sea ice melt and the resulting environmental conditions are putative key drivers of microbial eukaryotic community composition and bloom phenomenology. Our observations suggest that polar pelagic and ice-associated taxa (such as *Fragilariopsis cylindrus* or *Melosira arctica*) are relatively tolerant of more Atlantic oceanographic conditions. In contrast, temperate taxa (such as *Odontella aurita* or *Proboscia alata*) have limited potential to persist in colder ice-impacted waters. Thus, we hypothesize that sea-ice melt in the MIZ may hinder the northward expansion of temperate Atlantic taxa towards the CAO. This trend will continue even as Atlantic oceanographic conditions move further northwards.

## Supporting information

All supplementary figures and tables

## ACKNOWLEDGEMENTS

We thank Theresa Hargesheimer, Jana Bäger and Daniel Scholz for technical support of the RAS deployment. Moreover we thank the captains and crews of RV Polarstern for excellent support at sea, and the chief scientists for leading the various expeditions conducted for this study. Ship time for RV Polarstern was provided under grants AWI_PS99_00, AWI_PS100_01, AWI_PS107_05, AWI_PS114_01, AWI_PS121_01 of RV Polarstern. Our special thanks go to S. Neuhaus for bioinformatic support, K. Oetjen and S. Ziemann for excellent technical support in the laboratory, Eva-Maria Nöthig for critical reading of the manuscript, Martina Löbl for coordination of the FRAM-project.

## AUTHOR CONTRIBUTIONS

EO conducted the data analyses. WJvA, CB, MW, STV and KM are responsible for the sampling design. STV contributed nutrient data. WJvA contributed oceanographic data. EO, KM and OP interpreted the data, conceptualized the and drafted the manuscript. All authors contributed to improving the final manuscript, by contributions to the scientific interpretation of the data and the discussion of results.

## Data Availability Statement

The datasets generated during and/or analysed during the current study are available in the gitlab repository, https://gitlab.com/qtb-hhu/qtb-sda/framstrait_1718.

## ADDITIONAL INFORMATION

Correspondence and requests for materials should be addressed to Ellen Oldenburg.

