## Supplementary material for "Sea-ice melt determines seasonal phytoplankton dynamics and delimits the habitat of temperate Atlantic taxa as the Arctic Ocean atlantifies": All supplementary figures and tables

**Supplementary information**

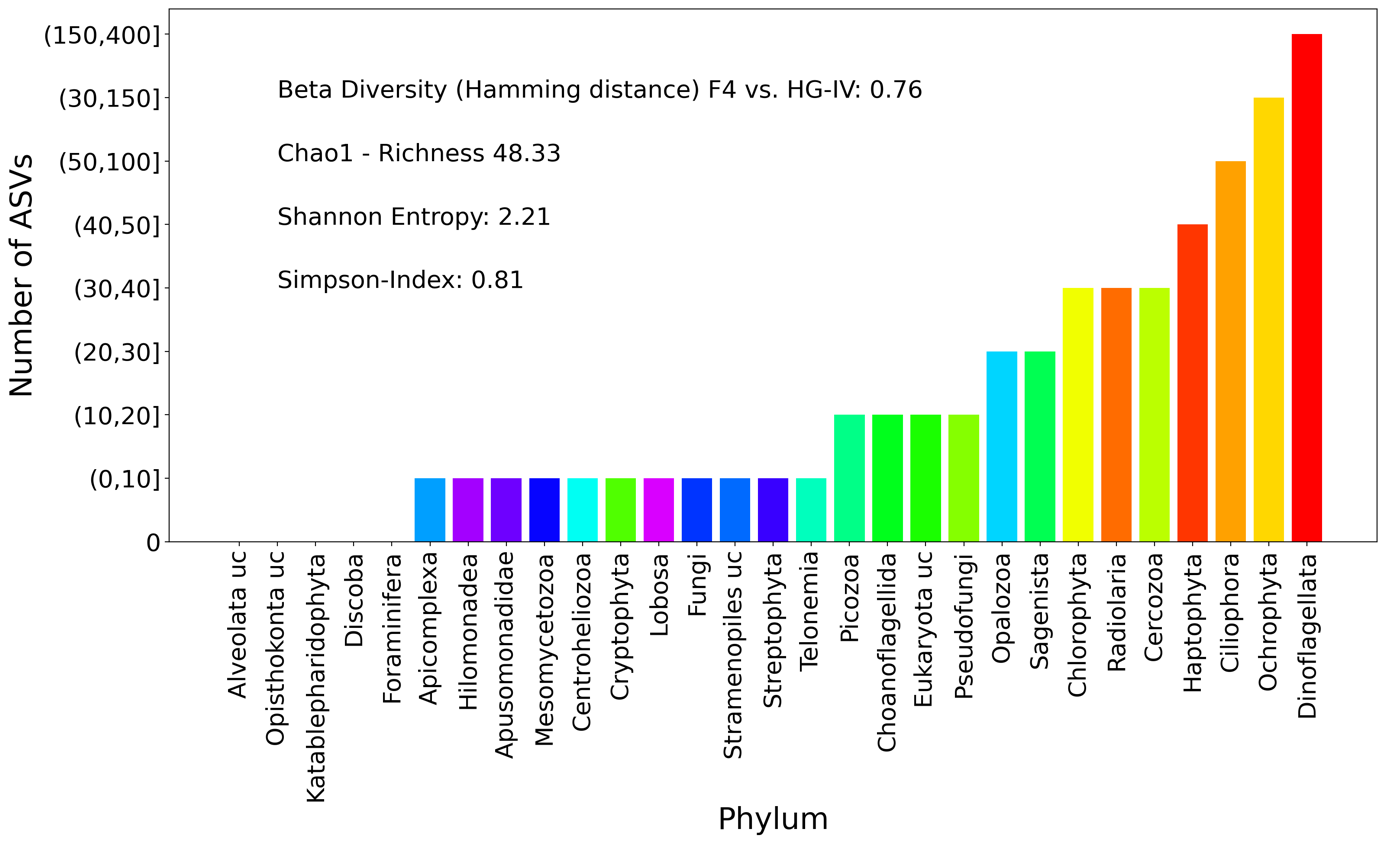

Figure S1: **Bio Diversity on Phylum Level:** The Bar plot show the number of ASV of the corresponding Phylum. Beta Diversity (Hamming distance) between both moorings and the alpha diversity for this mooring are displayed. The Richness by Chao1, the Shannon Entropy and the Simpson-Index as measurements for the alpha diversity are also calculated. **A**: HG-IV, **B**: F4.

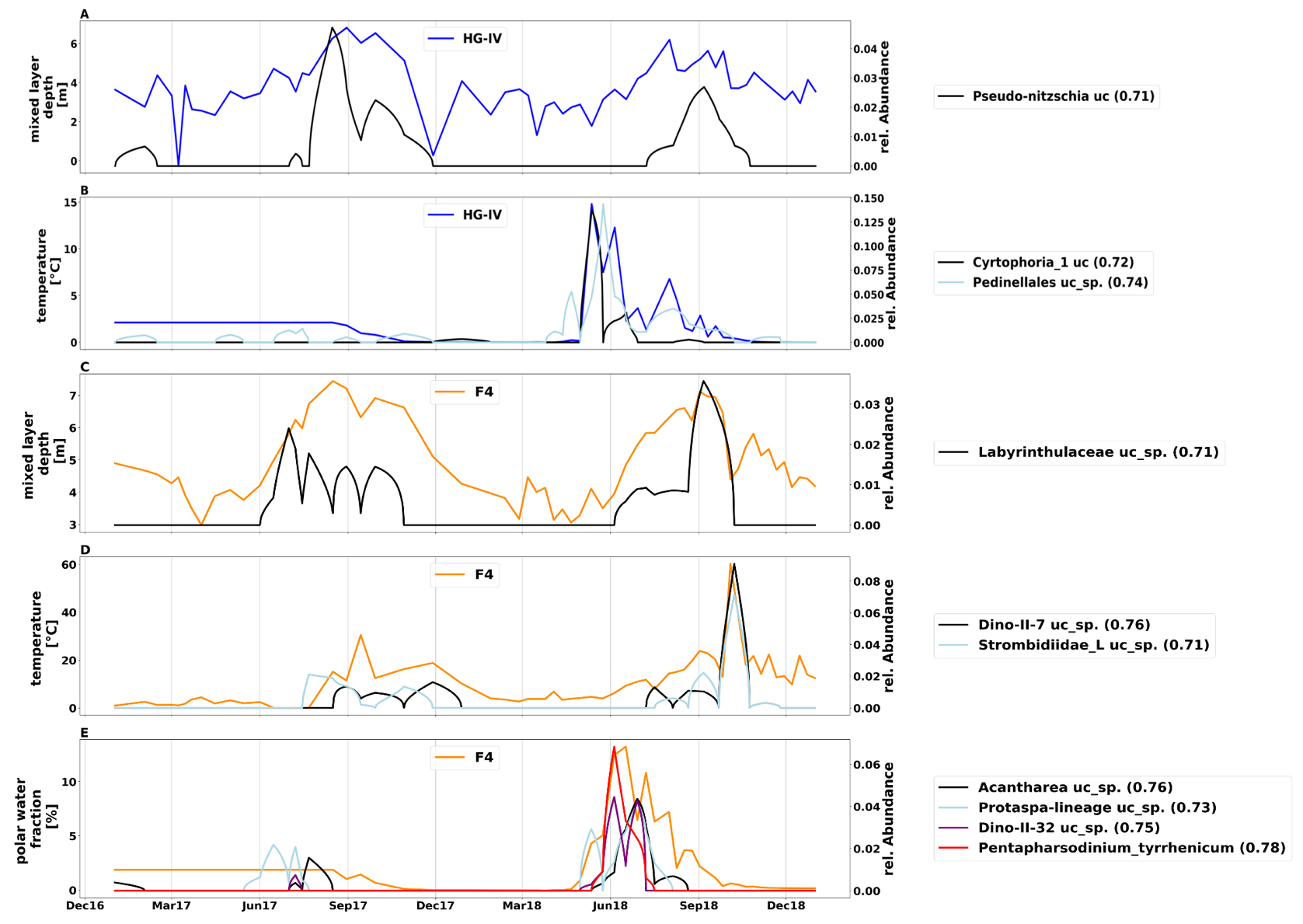

Figure S2: **Correlation between environment conditions and Species (ASVs):** Species with coefficient greater then 0.7, the value bracket indicate Pearson correlation coefficient are shown. **A**: HG-IV: mixed-layer depth, **B**: HG-IV: temperature, **C**: F4: mixed-layer depth, **D**: F4: temperature, **E**: F4: polar water fraction.

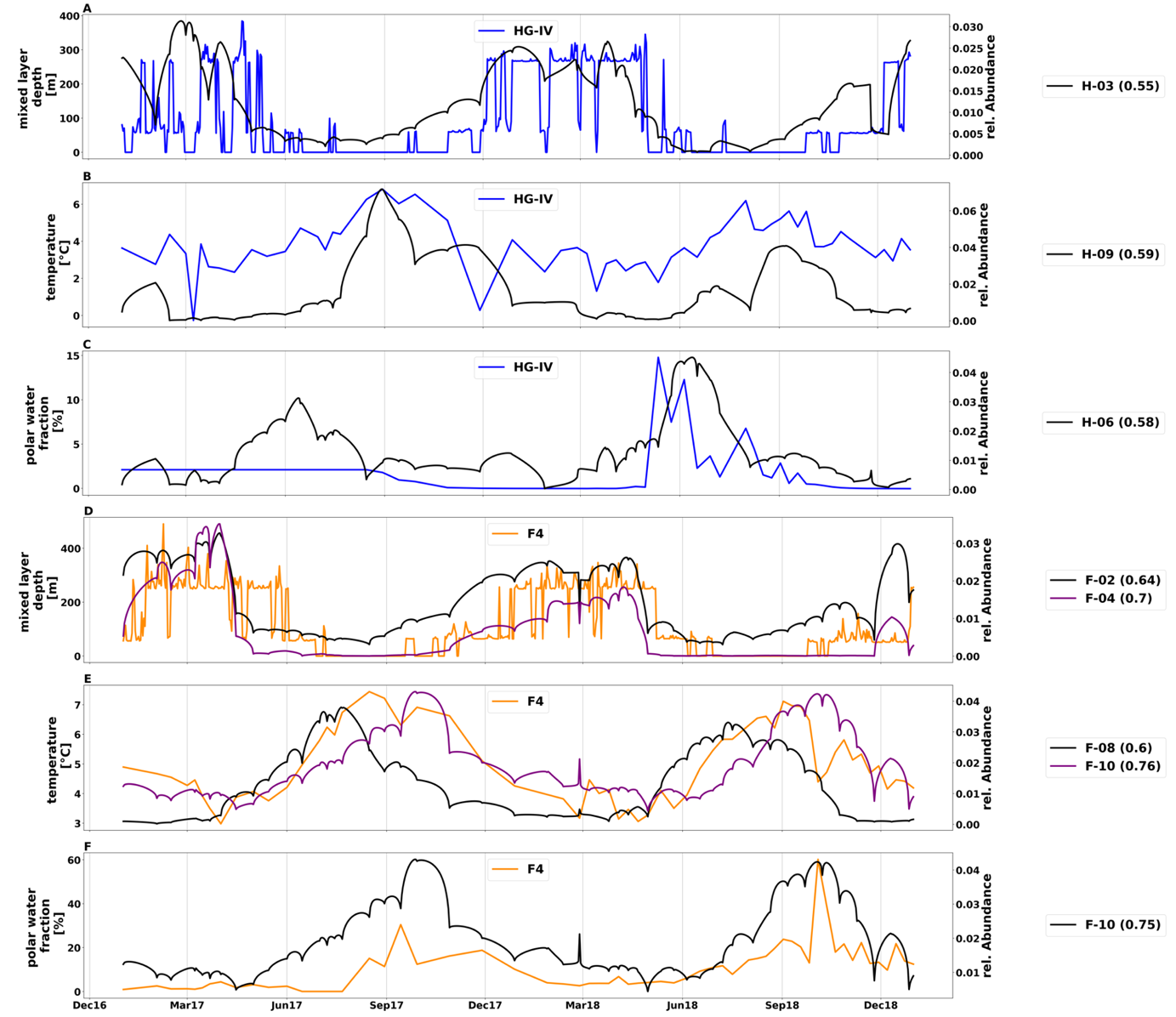

Figure S3: **Correlation between environment conditions and clusters:** Clusters with coefficient greater then 0.55, the value bracket indicate Pearson correlation coefficient are shown. **A**: HG-IV: mixed-layer depth, **B**: HG-IV: temperature, **C**: HG-IV: polar water fraction, **D**: F4: mixed-layer depth, **E**: F4: temperature, **F**: F4: polar water fraction.

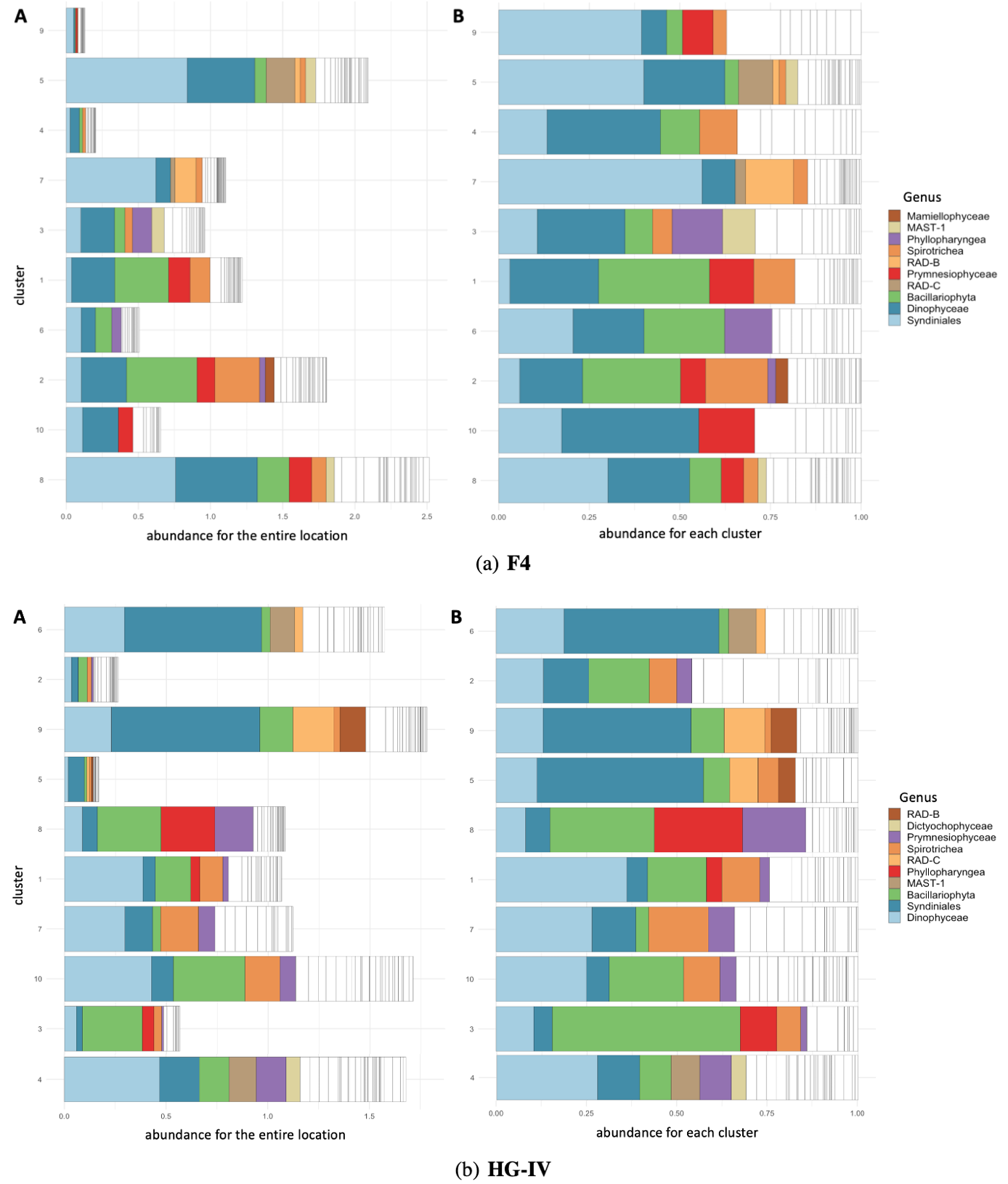

Figure S4: **Taxa bar-plot showing the top then microbial communities for both moorings.** Clusters are grouped according to their seasonality on the y-axis. Relative sequence abundances (%) of the ten most abundant eukaryotes classes, with all other genera grouped in "Other " (colored in white) are shown. The top 10 genera for HG-IV are shown ascending, starting with Dinophyceae (skyblue), Syndiniales (blue), Bacillariophyta (light green), MAST-1 (light brown), Phyllopharyngea (red), RAD-C (light orange), Spirotichea (orange), Prymnesiophyceae (purple), Dictyochophyceae (beach) and RAD-B (brown). The top 10 genera for F4 are shown ascending, starting with Syndiniales (skyblue), Dinophyceae (blue), Bacillariophyta (light green), RAD-C (light brown), Prymnesiophycaeae (red), RAD-B (light orange), Spirotichea (orange), Phyllopharyngea (purple), MAST-1 (beach) and Mamiellophyceae (brown). The x-axis indicates **A**: portion on the whole mooring for HG-IV, **B**: portion on the certain cluster for HG-IV., **C**: portion on the whole mooring for F4 and **D**: portion on the certain cluster for F4.

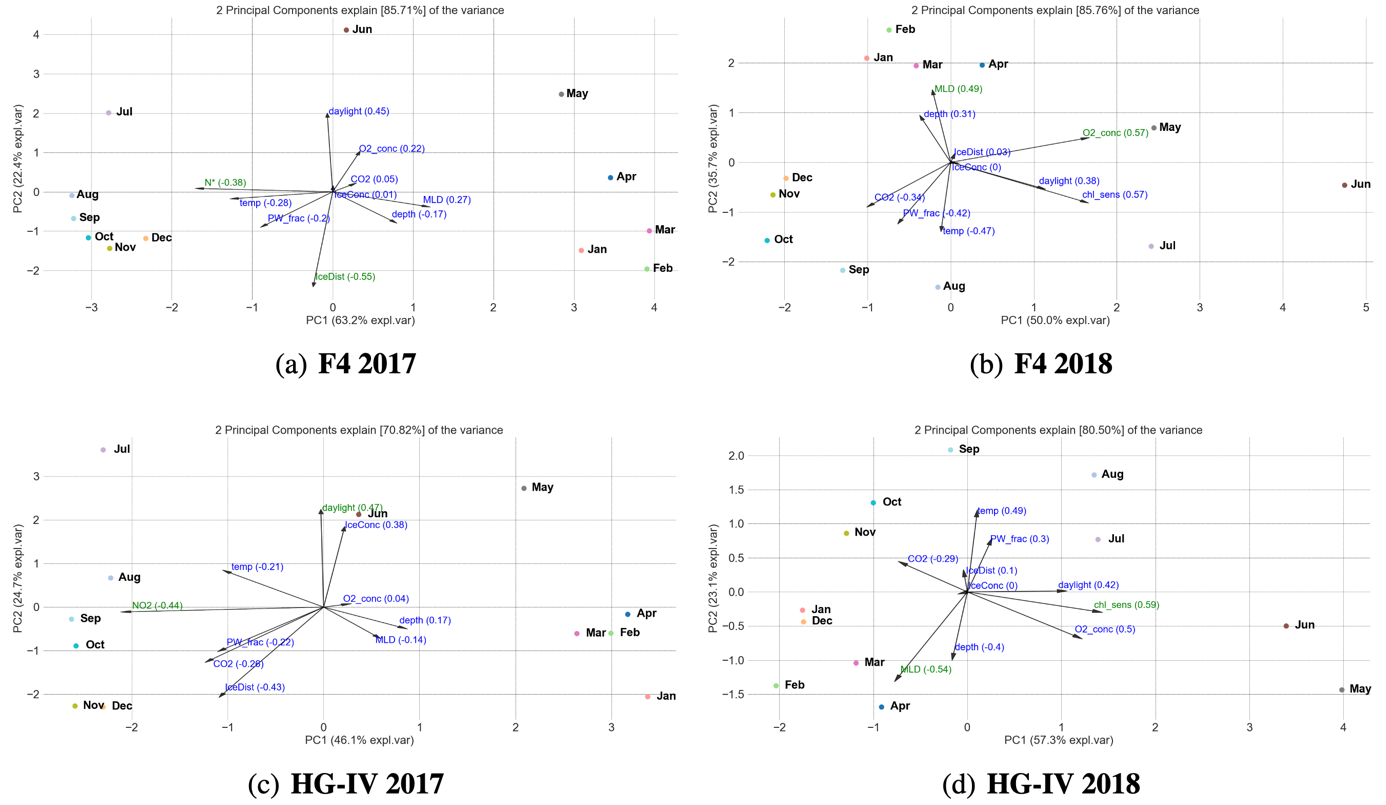

Figure S5: **Biplots for both mooring and both years**: First PCA component plotted versus the second PCA component of the abundance values for the ASV and the environment conditions on monthly bases aggregated. The arrows correspond to the most relevant features. The length of the arrow corresponds to the feature importance associated with this arrow. **A**: HG-IV 2017, **B**: HG-IV 2018, **C**: F4 2017 and **D**: F4 2018.

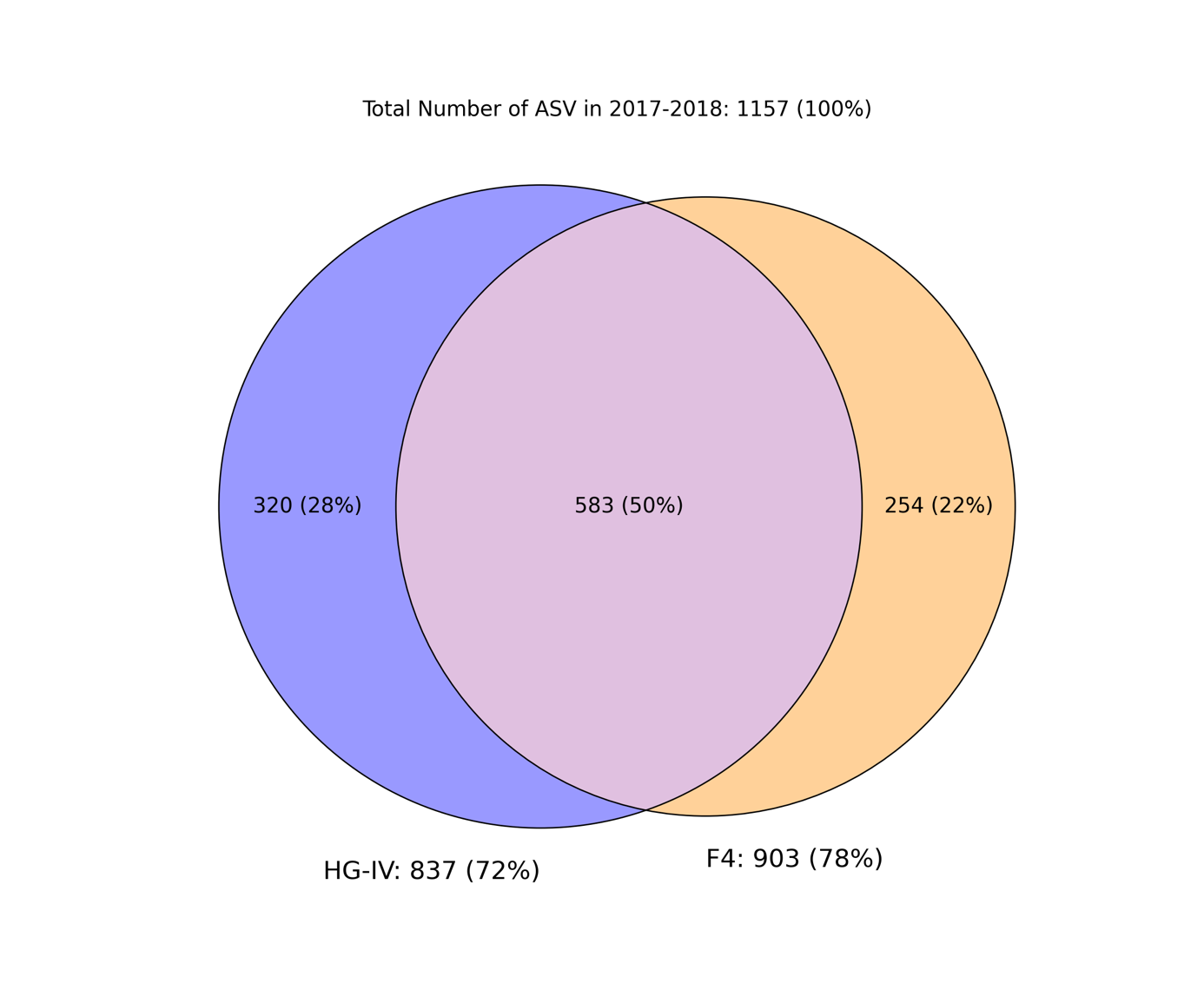

Figure S6: **Different ASV Communities for F4 and HG-IV:** Venn Diagram showing the Union (combination of cyan, dark orange and green), the Intersection (green) and the Set-Differences (F4 dark orange, HG-IV cyan) of the ASV for the F4 and HG-IV locations for 2017-2018.

Table S1: **Selected Species:** The species, the asv-name, the area under the curve 2017 (AUC17) and area under the curve 2018 (AUC18) and the quotients are shown.

| **Species** | **asv** | **Cluster** | **AUC17** | **AUC18** | **Quotient 17/18** | **Quotient 18/17** |
| --- | --- | --- | --- | --- | --- | --- |
| *Odontella_aurita* | euk_asv96 | H-05 | 3,2281 | 4.5659 | 0.707 | 1.4144 |
| *Fragilariopsis_cylindrus* | euk_asv207 | F-10 | 4.6039 | 4.3962 | 1.0472 | 0.9549 |
| *Fragilariopsis_sublineata* | euk_asv35 | F-08 | 23.7127 | 14.1505 | 1.6757 | 0.5967 |
| *Fragilariopsis_sublineata* | euk_asv35 | H-08 | 25.9609 | 13.7689 | 1.8855 | 0.5304 |
| *Pseudo-nitzschia_sp.* | euk_asv9 | F-10 | 31.291 | 30.224 | 1.0353 | 0.9659 |
| *Pseudo-nitzschia_sp.* | euk_asv9 | H-09 | 66.0964 | 42.8685 | 1.5418 | 0.6486 |
| *Melosira_arctica* | euk_asv244 | H-08 | 2.2228 | 0.247 | 8.9992 | 0.1111 |
| *Melosira_arctica* | euk_asv632 | H-02 | 5.2386 | 0 | ∞ | 0 |

| **cluster** | **max_peak_date_17** | **max_peak_17** | **max_peak_date_18** | **max_peak_18** |
| --- | --- | --- | --- | --- |
| H-01 | 20.02.17 | 0.03 | 26.11.18 | 0.059 |
| H-02 | 02.01.17 | 0.02 | 17.12.18 | 0.019 |
| H-03 | 25.02.17 | 0.031 | 31.12.18 | 0.027 |
| H-04 | 18.02.17 | 0.026 | 01.04.18 | 0.011 |
| H-05 | 20.04.17 | 0.033 | 25.04.18 | 0.044 |
| H-06 | 13.06.17 | 0.031 | 12.06.18 | 0.045 |
| H-07 | 26.05.17 | 0.036 | 29.07.18 | 0.048 |
| H-08 | 17.07.17 | 0.05 | 07.07.18 | 0.033 |
| H-09 | 29.08.17 | 0.072 | 07.09.18 | 0.041 |
| H-10 | 17.09.17 | 0.031 | 18.10.18 | 0.039 |
| F-01 | 29.10.17 | 0.007 | 27.10.18 | 0.052 |
| F-02 | 31.03.17 | 0.033 | 16.12.18 | 0.03 |
| F-03 | 12.02.17 | 0.033 | 26.01.18 | 0.023 |
| F-04 | 31.03.17 | 0.035 | 08.04.18 | 0.018 |
| F-05 | 19.03.17 | 0.03 | 29.07.18 | 0.029 |
| F-06 | 11.06.17 | 0.032 | 07.05.18 | 0.035 |
| F-07 | 09.09.17 | 0.033 | 12.04.18 | 0.024 |
| F-08 | 21.07.17 | 0.038 | 13.07.18 | 0.033 |
| F-09 | 10.09.17 | 0.026 | 24.11.18 | 0.055 |
| F-10 | 27.09.17 | 0.043 | 03.10.18 | 0.042 |

Table S2: **F4 and HG-IV Cluster max peaks per year:** For each mooring, for cluster and each year, the maximal peak date and the maximal peak are shown.

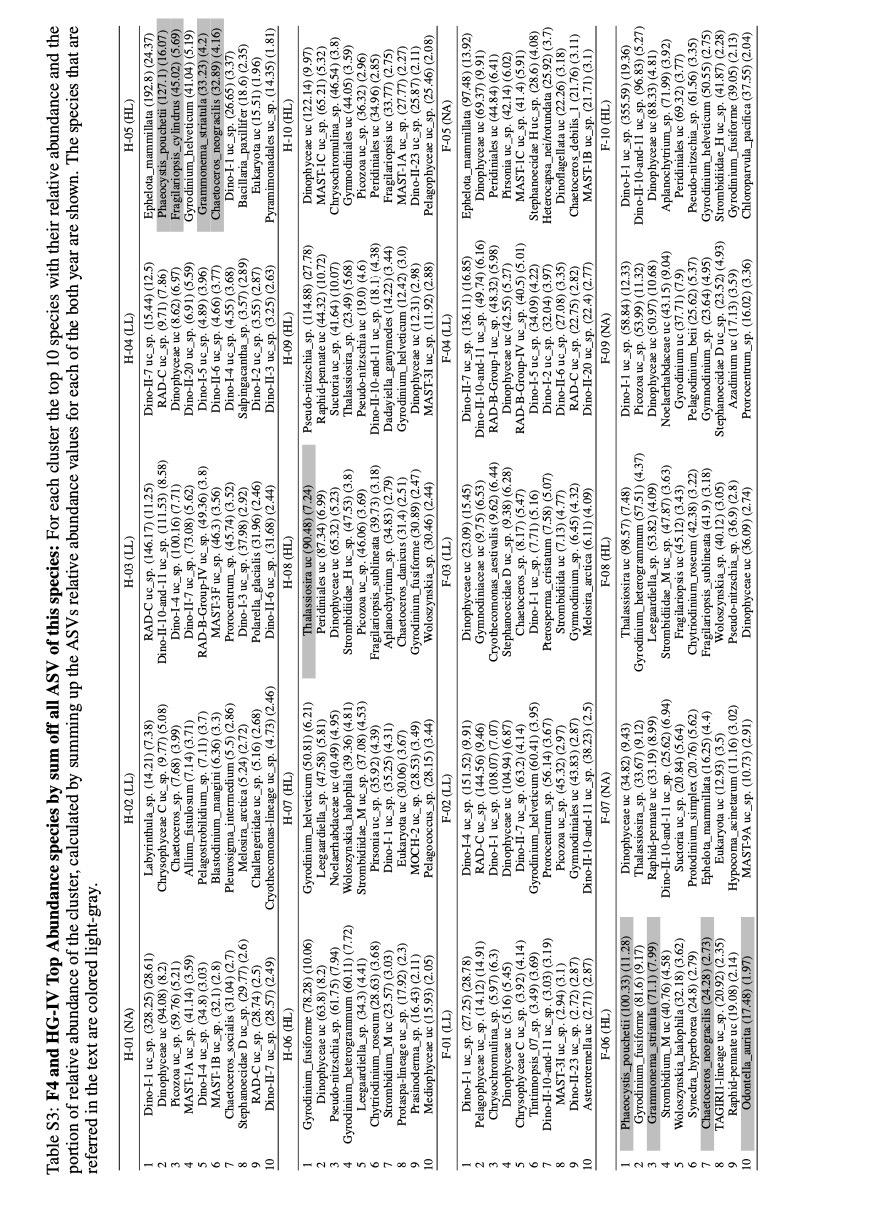

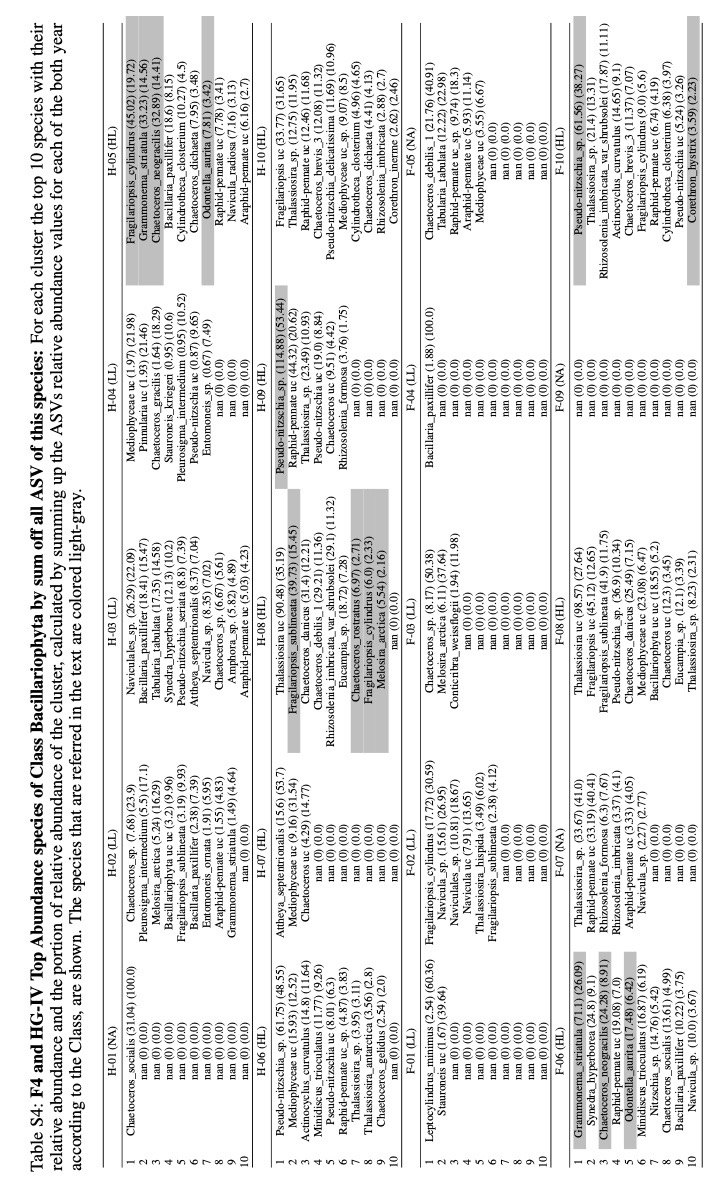

| **Name** | **Value** | **% shared ASV** | **% specific ASV** |
| --- | --- | --- | --- |
| Number of shared ASVs | 583 |  |  |
| Number of ASVs with non Zero Abundance | 559 |  |  |
| Number of MWR ASVs | 94 | 0.1612 | 0.5839 |
| Number of MLR ASVs | 67 | 0.1149 | 0.4161 |
|  |  | **STD** |  |
| q(F42017) | 1.234558 |  |  |
| q(HG2017) | 2.138047 |  |  |
| q(F42018) | 0.6025356 |  |  |
| q(HG2018) | 0.2653313 |  |  |
| median(p(F42017,HG-IV2017)) | 0.9245897 | 0.5691752 |  |
| median(p(F42018,HG-IV2018)) | 1.774752 | 5.317479 |  |
| median(t(F42017,HG-IV2017)) | 1.431774 | 4.379805 |  |
| median(t(F42018,HG-IV2018)) | 0.6382022 | 0.3910333 |  |
| **Kolmogorov-Smirnov test** | **D** | **p-value** |  |
| p(F42017,HG-IV2017) vs t(F42017,HG-IV2017) (two sided) | 0.3563036 | 6.25E-05 |  |
| p(F42017,HG-IV2017) vs t(F42017,HG-IV2017) (H_0 greater) | 0.3563036 | 3.13E-05 |  |
| p(F42018,HG-IV2018) vs t(F42018,HG-IV2018) (two sided) | 0.6592569 | 1.38E-14 |  |
| p(F42018,HG-IV2018) vs t(F42018,HG-IV2018) (H_0 less) | 0.6592569 | 1.37E-14 |  |
| **Ratios** |  |  |  |
| F4_2017_MWR_HP (polar) | 0.9245897 |  |  |
| F4_2018_MWR_HP | 0.45649254 |  |  |
| HG_2017_MWR_HP | 1 |  |  |
| HG_2018_MWR_HP | 0.31497868 |  |  |
| F4_2017_MLR_HP (atlant) | 0.66966442 |  |  |
| F4_2018_MLR_HP | 0.75761921 |  |  |
| HG_2017_MLR_HP | 0.46771657 |  |  |
| HG_2018_MLR_HP | 1.18711469 |  |  |
| quotient_of_mean_of_hg18_to_hg17 | 2.538107 |  |  |
| p(F42018,HG-IV2018) larger t(F42018,HG-IV2018) | 2.78086161 |  |  |
| t(F42017,HG-IV2017) larger p(F42017,HG-IV2017) | 1.54855067 |  |  |

Table S5: **Mixed Layer Regime ASVs are reduced 2017 in comparison to 2018 in HG-IV:** The charred ASVs of F4 and HG-IV were grouped into meltwater regimes (MWR) and mixed layer regimes (MLR) based on their abundance ratio. Quotients of the location wise ratios of MWR and MLR were calculated for each year. In addition, MWR and MLR ASV Abundance ratios were calculated for each location between the years. For each year, the groups MWR and MLR were then tested using the KS test, whose distribution was greater/smaller using a one-sided test. These eight quotients were used to calculate the pairwise ratio of the MRW and MLW on different locations and years and normalized to 1.
